# UP-DOWN cortical dynamics reflect state transitions in a bistable balanced network

**DOI:** 10.1101/083626

**Authors:** Daniel Jercog, Alex Roxin, Peter Barthó, Artur Luczak, Albert Compte, Jaime de la Rocha

## Abstract

In the idling brain, neuronal circuits often exhibit transitions between periods of sustained firing (UP state) and quiescence (DOWN state). Although these dynamics occur across multiple areas and behavioral conditions, the underlying mechanisms remain unclear. Here we analyze spontaneous population activity from the somatosensory cortex of urethane-anesthetized rats. We find that UP and DOWN periods are variable (i.e. non-rhythmic) and that the population rate shows no significant decay during UP periods. We build a network model of excitatory (E) and inhibitory (I) neurons that exhibits a new bistability between a quiescent state and a balanced state of arbitrarily low rate. Fluctuating inputs trigger state transitions. Adaptation in E cells paradoxically causes a marginal decay of E-rate but a marked decay of I-rate, a signature of balanced bistability that we validate experimentally. Our findings provide evidence of a bistable balanced network that exhibits non-rhythmic state transitions when the brain rests.

## Introduction

A ubiquitous pattern of spontaneous cortical activity during synchronized brain states consists in the alternation between periods of tonic firing (UP states) and periods of quiescence (DOWN states) (Luczak et al., 2007; Steriade et al., 1993a; Timofeev et al., 2001). Cortical UP and DOWN dynamics take place during slow-wave-sleep (SWS) (Steriade et al., 1993a) and can also be induced by a number of anesthetics (Steriade et al., 1993a). More recently however, similar synchronous cortical dynamics have been observed not only in awake rodents during quiescence (Luczak et al., 2007; Petersen et al., 2003), but also in animals performing a perceptual task, both rodents (Sachidhanandam et al., 2013; Vyazovskiy et al., 2011) and monkeys (Engel et al., 2013).

Spontaneous activity resembling UP and DOWN states has been found in cortical slices *in vitro* (Cossart et al., 2003; Fanselow and Connors, 2010; Mann et al., 2009; Sanchez-Vives and McCormick, 2000), in slabs (Timofeev et al., 2000) and *in vivo* under extensive thalamic lesions (Steriade et al., 1993b). This has led to suggest that the underlying mechanism had an intracortical origin. In such scenario, the standard hypothesis postulates that during UP periods a fatigue cellular mechanism – e.g. spike frequency adaptation or synaptic short-term depression – decreases network excitability until the state of tonic firing can no longer be sustained and the cortical network switches into a DOWN state (Contreras et al., 1996; Sanchez-Vives and McCormick, 2000). During DOWN periods, in the absence of firing, the fatigue variables recover until the circuit becomes self-excitable and autonomously transitions into an UP state (Cunningham et al., 2006; Le Bon-Jego and Yuste, 2007; Poskanzer and Yuste, 2011; Sanchez-Vives and McCormick, 2000; Timofeev et al., 2000). This mechanism of activity dependent negative feedback causing oscillatory UP-DOWN dynamics has been implemented by several computational models (Bazhenov et al., 2002; Benita et al., 2012; Chen et al., 2012; Compte et al., 2003b; Hill and Tononi, 2005; Parga and Abbott, 2007). However, although commonly described as a slow oscillation, the rhythmicity of UP-DOWN dynamics has not been systematically quantified and seems to depend on the details of the preparation (Chauvette et al., 2011; Erchova et al., 2002; Lampl et al., 1999; Ruiz-Mejias et al., 2011).

Alternatively, there is strong evidence suggesting that UP-DOWN transitions in neocortical circuits are coupled with activity in other areas. Thalamocortical neurons for instance can endogenously oscillate at low frequencies (Hughes et al., 2002; McCormick and Pape, 1990), cause cortical UP states when stimulated (Rigas and Castro-Alamancos, 2007) or modulate the UP-DOWN dynamics when suppressed (David et al., 2013; Lemieux et al., 2014) and their spontaneous activity correlates with UP state onset (Contreras and Steriade, 1995; Slézia et al., 2011; Ushimaru et al., 2012). Moreover, the timing of hippocampal sharp-wave ripples (Battaglia et al., 2004), or basal ganglia activity (Ushimaru et al., 2012) also tends to precede DOWN to UP transitions. Finally, intracortical stimulation can effectively cause UP-DOWN transitions (Beltramo et al., 2013; Shu et al., 2003) even when only a few dozen neurons are stimulated (Stroh et al., 2013). In total, these findings describe a system whose macroscopic UP-DOWN dynamics are sensitive to temporal fluctuations of both external inputs and local circuit activity. Such a network would in principle generate unpredictable and therefore irregular UP-DOWN dynamics, since transitions are no longer dependent exclusively on local cortical internal dynamics.

The interplay of fatigue and fluctuations causing transitions between two states has been theoretically studied in the developing spinal cord (Tabak et al., 2011, 2000), and in the context of UP-DOWN dynamics mostly in excitatory networks (Holcman and Tsodyks, 2006; Lim and Rinzel, 2010; Mattia and Sanchez-Vives, 2012; Mejias et al., 2010). Models of spontaneous activity are however theoretically founded on the balance between excitatory (E) and inhibitory (I) populations (Amit and Brunel, 1997; van Vreeswijk and Sompolinsky, 1998), a dynamic state that can quantitatively mimic population spiking activity during desynchronized states (Renart et al., 2010). In spite of this, there is still no simple EI network model that, building on a balanced state, can exhibit bistability between a low-rate and a quiescent state (Latham et al., 2000). To develop such a model, we first performed population recordings of ongoing cortical activity during synchronized brain state epochs in rats under urethane anesthesia (Détári and Vanderwolf, 1987; Luczak et al., 2007; Murakami et al., 2005; Whitten et al., 2009). Analysis of multi single-unit spiking dynamics, showed irregular UP and DOWN periods and no decay of the average rate during UPs. Given these constraints, we built an EI network model that, capitalizing on the firing threshold non-linearity and the asymmetry of the E and I transfer functions, exhibited a novel type of bistability with a quiescence (DOWN) and a low-rate state (UP). External input fluctuations into the network caused the irregular UP-DOWN transitions. Adaptation in E cells in contrast, did not cause transitions and had a different effect on the E rate in each of the two states: while it exhibited recovery during DOWN periods, it showed almost no decay during UP periods due to the balanced nature of the UP dynamics. Our model provides the first EI network that exhibits stochastic transitions between a silent and a balanced attractor matching the statistics of UP and DOWN periods and population rate time-courses observed in the cortex.

## Results

To investigate the mechanisms underlying the generation of spontaneous cortical activity, we recorded the spiking activity from large populations of neurons (mean±SD = 64±23 cells) in deep layers of somatosensory cortex of urethane-anesthetized rats (n=7 animals) (Barthó et al., 2004; Luczak et al., 2009). Because brain state under urethane can vary spontaneously (Détári and Vanderwolf, 1987; Luczak et al., 2007; Murakami et al., 2005; Whitten et al., 2009), we selected the most clearly synchronized epochs characterized by the stable presence of high-amplitude, slow fluctuations in cortical local field potential (LFP) signals (Fig. 1A; see Methods) (Harris and Thiele, 2011; Steriade et al., 2001). During these epochs, the instantaneous population rate *R*(*t*), defined from the merge of all the recorded individual spike trains, displayed alternations between periods of tonic firing and periods of silence (Luczak et al., 2007), a signature of UP and DOWN states from an extracellular standpoint (Fig. 1B-C) (Cowan and Wilson, 1994; Sanchez-Vives and McCormick, 2000; Steriade et al., 1993a). Despite the clear presence of UP and DOWN states, the population activity displayed no clear traces of rhythmicity as revealed by strongly damped oscillatory structure in both autocorrelograms of LFP and *R(t)* (Fig 1D and 1E, respectively). Motivated by this, we hypothesized that the cortical circuit might transition between two network states in a random manner (Deco et al., 2009; Mejias et al., 2010; Mochol et al., 2015). Using a probabilistic hidden semi-Markov model (Chen et al., 2009), we inferred the instantaneous state of the circuit from the population rate *R(t)* by extracting the sequence of putative UP (U) and DOWN (D) periods (Fig. 1C, Methods).

**Figure 1.**
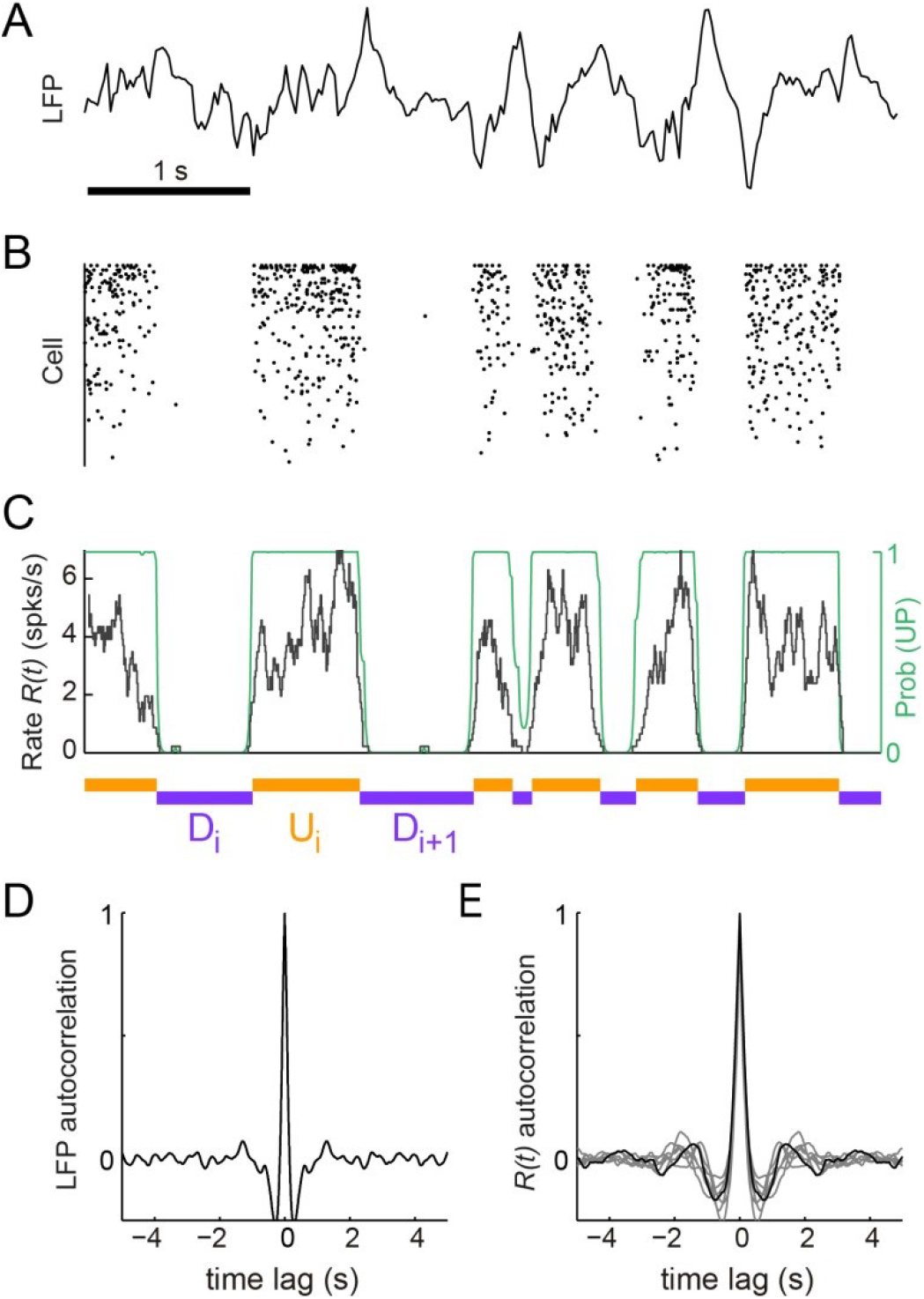
*Synchronized brain activity under urethane anesthesia in the rat somatosensory cortex and the detection of putative UP and DOWN periods.* **(A).** Local field potential during 5 s of synchronized state displaying high-amplitude slow fluctuations. **(B)** Population raster of 92 simultaneously recorded single units exhibiting the alternation between periods of tonic spiking activity and periods of neural quiescence (cells sorted based on mean firing rate). **(C)** Instantaneous population rate *R(t)* (grey) is used to identify putative U (orange) and D (purple) periods. The detection algorithm is based on fitting a Hidden Markov Model (HMM) and computing the posterior probability of the hidden state being in an UP state (green) (see methods). **(D)** Average autocorrelogram of LFP (20-s windows) for one example experiment. **(E)** Average autocorrelogram of *R(t)* for different (n=7) experiments (example experiment in black).

### UP and DOWN duration statistics during synchronized states

The statistics of U and D period durations showed skewed gamma-like distributions (Fig. 2A and 2B right; Supp. Fig 1). The mean duration across different experiments displayed a wide range of values (Fig. 2B left; mean±SD: <U>=0.43±0.19 s, <D>=0.46±0.1 s, n=7), whereas the coefficients of variation CV(U) and CV(D) of U and D periods, defined as the standard deviation divided by the mean of the period durations within experiments, were systematically high (Fig. 2B middle, mean±SD: CV(U)=0.69±0.09, CV(D)=0.69±0.1). The irregularity in the U and D periods did not result from slow drifts in the mean U or D durations caused by variations of brain state as confirmed by computing the CV_2_ (Holt et al., 1996), a local measure of irregularity that is less affected by slow variations in the statistics (mean±SD: CV_2_(U)=0.86±0.13, CV_2_(D)=0.75±0.17; see Methods). The high variability of U and D periods is consistent with the non-periodicity of the dynamics revealed in the autocorrelation function (Fig. 1D-E).

**Figure 2.**
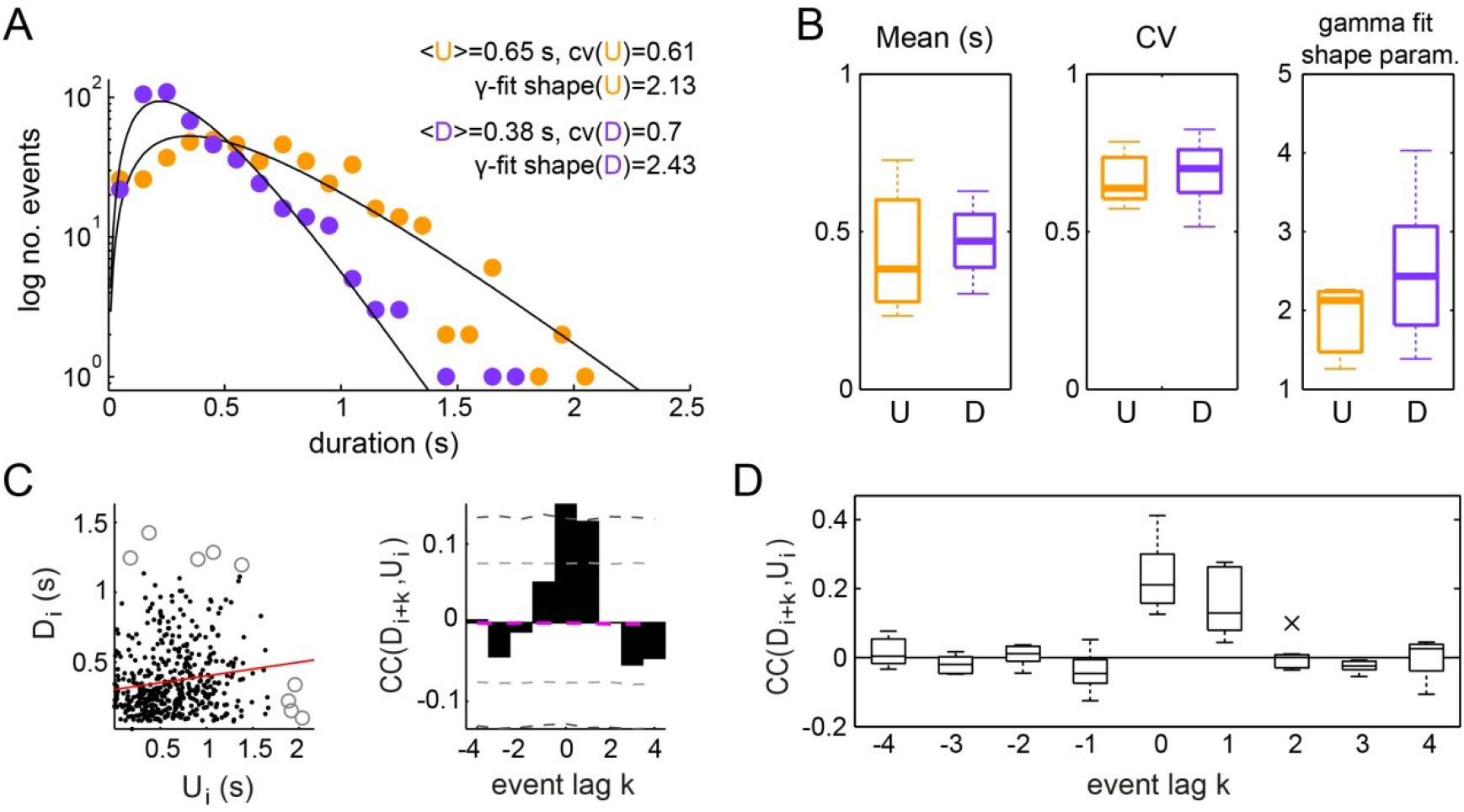
*Statistics of U and D periods during synchronized brain activity.* **(A)** Distribution of U and D durations for one example experiment (same as Fig 1). Inset shows the mean and coefficient of variation (CV) of U and D durations. **(B)** Summary of period duration mean (left) and CV (right) across experiments (n=7 rats). While average durations are quite heterogeneous across experiments, the period duration variability is consistently large. **(C)** Left: D duration (D_i_) vs the consecutive U duration (U_i_) exhibit weak but significant serial correlation (circle marks showing values away than 3 standard deviations of the mean, were discarded for correlation analysis; red line showing linear regression). Right: Cross-correlogram between the D_i_ and U_i_ sequences for different lags (k) in a single experiment. Magenta dashed line represent the mean cross-correlogram from a local shuffled (see Methods). Light (dark) grey dashed line showing 95% C.I. point-wise (global) error bands. **(D)** Summary of cross-correlation analysis for the different experiments, displaying consistent positive correlations across experiments for lags k=0 and k=1.

We then asked whether the lengths of U and D periods were independent, as if the transitions between the two network states would reset the circuit’s memory, or if in contrast they were correlated by a process impacting the variability of several consecutive periods. We computed the linear cross-correlation *Corr*(*U*_*i*_*,D*_*i+k*_) (Fig. 2C left, for *k*=0) between pairs of periods separated in the D-U sequence by a lag *k* (Fig. 2C, right). The cross-correlation *Corr(U*_*i*_*, D*_*i +k*_) displayed consistently nonzero values for k=0 and k=1 (mean±SD: 0.21±0.09, 0.17±0.09, respectively; significant cross-correlation in 6/7 animals, *P*<0.05 permutation test), whereas remained close to zero for the rest of lags, showing that period duration relationship is limited to adjacent periods (Fig 2C-D). The positive correlation between adjacent periods was not due to slow changes in their duration, as we corrected by the correlation obtained from surrogate D-U sequences obtained from shuffling the original sequence within 30 second windows (see Methods). Positive correlations between consecutive periods of activity and silence can be generated when fluctuation driven transitions are combined with an adaptive process such as activity-dependent adaptation currents (Lim and Rinzel, 2010; Tabak et al., 2000): if a fluctuation terminates a U period prematurely without much build-up in adaptation, the consecutive D period also tends to be shorter as there is little adaptation to recover from. However, a major role of adaptation currents in dictating UP-DOWN dynamics (Compte et al., 2003b) seems at odds with the lack of rhythmicity and the highly variable U and D durations, indicative of a stochastic mechanism causing the transitions between network states.

### Spiking activity during UP and DOWN states

We next searched for more direct evidence of an adaptive process by examining the time course of the population firing rate *R(t)* during U and D periods (see Fig. 1C; see Methods). The mean firing rate in U periods was low (mean±SD: 3.72±0.9 spikes/s, n=7). Moreover, D periods displayed occasional spiking (mean±SD rate 0.018±0.007 spikes/s; see e.g. Fig. 3A-B and Supp. Fig. S2), in contrast with the idea that DOWN periods do not display spiking activity (Chauvette et al., 2010), but see (Compte et al., 2003b). Thus, our hypothesis was that adaptation currents, if present, would induce a decay in *R(t)* during Us and an increase during Ds, and this impact on *R(t)* dynamics should be more evident during longer periods due to a larger accumulation (during Us) or recovery (during Ds) of the adaptation. For each experiment, we aligned the rate *R(t)* at the DOWN-to-UP (DU) and UP-to-DOWN (UD) transition times (Fig. 3A). We then computed the average rates *R*_*DU*_(*τ*) and *R*_*UD*_(*τ*) across all DU and UD transitions, respectively, with *τ* = 0 representing the transition time (Fig. 3B; mean across experiments = 598 transitions; range 472-768). Because Us and Ds had different durations, we selected long periods (U, D > 0.5 s) and compared *R*_*DU*_(*τ*) and *R*_*UD*_(*τ*) at the beginning and end of each period (mean number of Us 181, range 61-307; Ds 202, range 55-331). To specifically assess a change in rate during the U period, we compared the average *R*_*DU*_(*τ*) in the time window *τ*= (50, 200) ms (beginning of U) with the average *R*_*UD*_(*τ*) in the window *τ* = (-200, -50) ms (end of U), which we referred to as U-onset and U-offset windows, respectively. The windows were chosen 50 ms away from τ= 0 to avoid the transient change due to the state transitions (Fig. 3C-D). We found no significant mean difference between population average rate at U-onset and U-offset windows across our experiments (mean±SD onset minus offset population rate 0.04 ± 0.40 spikes/s, *P=1*, Wilcoxon signed rank, n=7 animals). The equivalent analysis performed on D periods yielded a small but significant mean increase in the population rate between the D-onset and D-offset windows (mean±SD −0.014 ± 0.013 spikes/s, *P*=0.047, Wilcoxon signed rank test). To examine in more detail the lack of population rate change during Us, we looked at the modulation of individual neuron rates normalized by the overall temporal average of each unit (Fig. 3E). We found that the change between U-onset and U-offset averaged across all our neurons (n=448 cells) was not significantly different from zero (Fig. 3E right, mean±SD of the onset vs offset difference of normalized rates 0.057 ± 1.163, *P*=0.12, Wilcoxon signed rank test) but that the recovery during D periods was significant (Fig. 3E left; mean±SD −0.015 ± 0.087, *P*=0.0002, Wilcoxon signed rank test). Some individual neurons however did show a significant modulation between U-onset and U-offset, but the decrease found in a fraction of the neurons was compensated with a comparable increase in another fraction of neurons (Fig. 3E right). Thus, at the population level, spiking activity during U periods displayed a sustained time course with no significant traces of rate adaptation.

**Figure 3.**
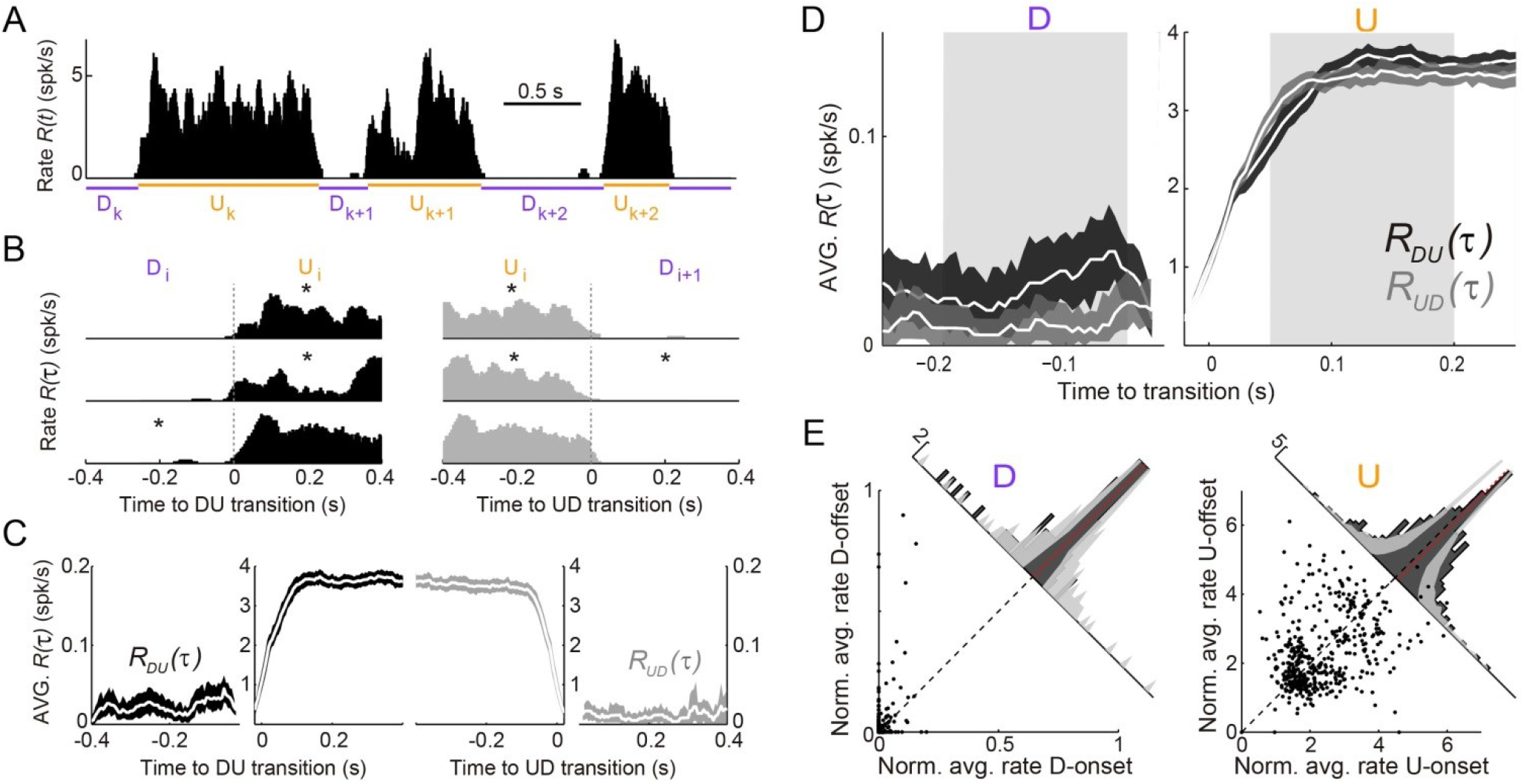
*Population spiking statistics during U and D periods.* **(A)** Example of instantaneous population rate R(t) with U and D detected periods (as in Fig.1). **(B-C)** Each U period is aligned at the DU (B, left) and UD (B, right) transition times in order to compute the instantaneous population rate averaged across transitions *R*_*DU*_(*T*) (C, dark grey) and *R*_*UD*_(*T*) (C, light grey), respectively. Only periods longer than 0.5 s (asterisks in B) were included in the average. **(D)** Comparison of population rate at the onset and offset of Us and Ds done by overlaying *R*_*DU*_(*t*) and a time-reversed *R*_*ud*_(*t*). Onset and offset windows defined during D and U periods (shaded) were used to test significance of changes in the rate. **(E)** Normalized firing rates from all individual neurons (448 cells from n=7 animals) during onset and offset windows. Left: D periods. Right: U periods. Average across cells is shown in red. Gray bands show 95#x0025; C.I. of the histograms obtained from onset-offset shuffled data (see Methods).

### Rate model for UP and DOWN dynamics

To understand the network and cellular mechanisms underlying the generation of stochastic U-D dynamics, showing U-D serial correlations and sustained rates during U periods, we analyzed a computational rate model composed of an excitatory (E) population recurrently coupled with an inhibitory (I) population (Latham et al., 2000; Ozeki et al., 2009; Tsodyks et al., 1997; Wilson and Cowan, 1972). The excitatory-inhibitory (EI) network model described the dynamics of the mean instantaneous rates *r*_*E*_ and *r*_*l*_ of each population in the presence of fluctuating external inputs. In addition, the E population included an adaptation mechanism, an additive hyperpolarizing current *a* that grew linearly with the rate *r*_*E*_ (Fig. 4A; see Methods). We did not consider adaptation in the inhibitory population for simplicity, and because inhibitory neurons show little or no spike-frequency adaptation when depolarized with injected current (McCormick et al., 1985). Our aim was to search for a regime in which, in the absence of adaptation and external input fluctuations, the network exhibited bistability between a quiescent (D) and a low-rate state (U) fixed point. Although bistability in low-dimensional EI networks has been described since the seminal work of Wilson & Cowan (1972), previous models primarily sought to explain bistability between a low-rate and a high-rate state, and exploited the combination of expansive and contractive non-linearities produced by the transfer function (Amit and Brunel, 1997; Renart et al., 2007; Wilson and Cowan, 1972), short-term synaptic plasticity (Hansel and Mato, 2013; Mongillo et al., 2008) or the divisive effect inhibitory conductances (Latham et al., 2000) (see Discussion). We found that bistability between D and U states can be robustly obtained solely using the expansive nonlinearity of the transfer function caused by the spiking threshold. Given this, we choose the simplest possible transfer function with a threshold: a threshold-linear function (Fig.4B, see Methods). Our choice to only use an expansive threshold non-linearity constrained strongly the way in which the network could exhibit bistability as can be deduced by plotting the nullclines of the rates *r*_*E*_ and *r*_*I*_ (Fig. 4C): only when the I nullcline was shifted to the right and had a larger slope than the E nullcline, the system exhibited two stable attractors (Eq. 20 in Methods). This configuration of the nullclines was readily obtained by setting the threshold and the gain of the I transfer function larger than those of the E transfer function (Fig. 4B), a distinctive feature previously reported when intracellularly characterizing the *f - I* curve of pyramidal and fast spiking interneurons in the absence of background synaptic activity (Cruikshank et al., 2007; Schiff and Reyes, 2012). This novel bistable regime yielded a quiescent D state, and arbitrarily low firing rates for both E and I populations during U states, depending on the values of the thresholds and the synaptic weights (Fig. 4C). This is remarkable as in most bistable network models the rate of the sustained activity state is constrained to be above certain lower bound (see Discussion). Moreover, in this bistable regime, the U state is an inhibition-stabilized state, a network dynamical condition in which the excitatory feedback is so strong that would alone be unstable, but is balanced with fast and strong inhibitory feedback to maintain the rates stable (Ozeki et al., 2009; Tsodyks et al., 1997) (see Methods).

**Figure 4.**
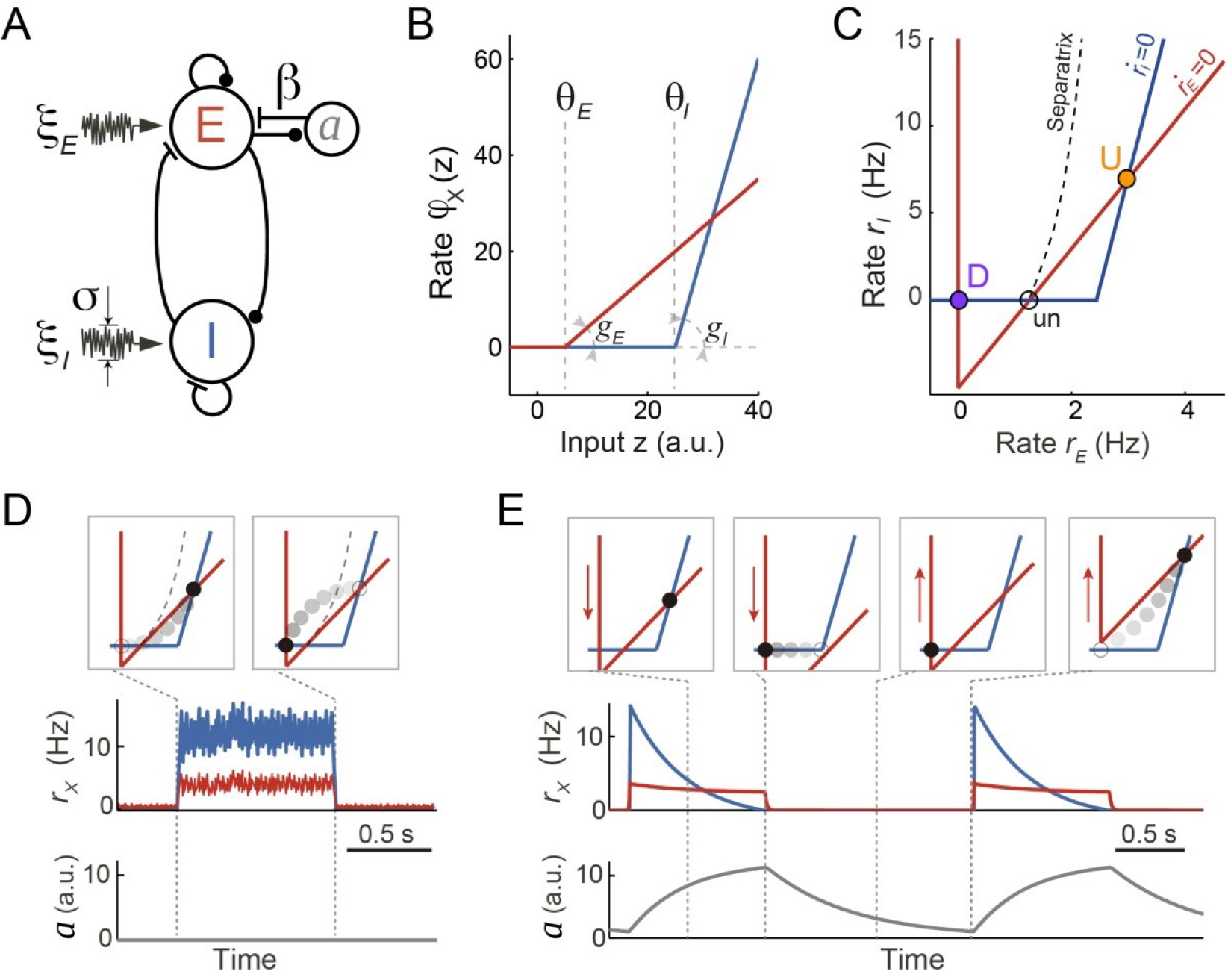
*Rate model for fluctuations and adaptation induced UP and DOWN dynamics.* **(A)** Network composed of recurrently connected inhibitory (I, blue) and excitatory (E, red) populations, with E exhibiting rate adaptation *a(t)* and both populations receiving independent fluctuating external inputs. **(B)** Transfer functions for the E and I populations are threshold-linear with unequal thresholds θ_e_ < θ_*l*_ and unequal gains g_e_ < g_i_. This marked asymmetry is at the origin of the bistability obtained in the network. **(C)** In the absence of adaptation, the phase plane of rates *r*_e_ vs. *r*_i_ shows the E and I nullclines (red and blue, respectively) whose intersections determine two stable (U and D) and one unstable (un) fixed points. The separatrix (dashed line) divides the phase plane into the basins of attraction of the D and U stable points. **(D, E)** Schematics of fluctuations-induced DU and UD transitions in the absence of adaptation (β=0) and adaptation-induced transitions in the absence of fluctuations (σ=0), respectively. Traces of *r*_*e*_*(t)*, *r*_*l*_*(t)* and adaptation *a(t)* illustrate steady fluctuating rates during U periods when there is no adaptation (D), and a periodic alternation between U and D characterized by a strongly decaying I rate during Us when there is no fluctuations (E). Top insets show the network trajectories in the phase-plane taken at different time points (vertical dotted lines). Notice the downward (upward) displacement of the E-nullcline during U (D) periods (red arrows in E). Connectivity parameters: *J*_*EE*_ = 5, *J*_*EI*_ = 1, *J*_*IE*_ = 10, *J*_*II*_ = 0.5 s; Transfer function parameters: *g*_*E*_ = 1, *g*_*I*_ = 4 Hz, θ_*E*_ = 0, θ_I_ = 25 a.u.

There are two ways in which transitions between U and D states can occur. On the one hand, transitions could be driven by external input fluctuations, which were modeled as a stochastic process with zero mean and short time constant (Fig. 4D). This fluctuating input reflected either afferents coming from other brain areas whose neuronal activity was stochastic and uncorrelated with the cortical circuit internal dynamics or the stochasticity of the spiking happening during U periods which was not captured by the dynamics of the rates (Holcman and Tsodyks, 2006; Lim and Rinzel, 2010). On the other hand, in the absence of fluctuations, state transitions could also occur solely driven by adaptation currents (Fig. 4E). Because the adaptation time constant was much longer than the time constants of the E and I rates, the dynamics of the rates *r*_*E*_*(t)* and *rI(t)* relaxing rapidly to their steady-state can be decoupled from the slow changes in *a(t)* (Latham et al., 2000; Rinzel and Lee, 1987). The network dynamics can be described in the phase plane (*r*_*E*_*(t)*, *r_I_(t)*) with variations in a(t) causing a displacement of the E-nullcline. In particular, during U periods the build-up in adaptation produced a downward displacement of the E-nullcline (Fig. 4E). If adaptation strength β was sufficiently large the displacement increased until the U state was no longer a fixed point and the network transitioned to the only stable fixed point D. Recovery of adaptation during D periods shifted the E-nullcline upwards until the D state disappeared and there was a transition to the U state (Fig 4E). In this limit cycle regime the network exhibited an oscillatory behavior with a frequency close to the inverse of the adaptation recovery time constant. When the two types of transitions are combined, two types of stability in U and D states can be distinguished: (1) metastable, referred to a state that was stable to the dynamics of both the rates and the adaptation but could transition away due to input fluctuations; (2) quasi-stable, referred to a state that was stable for the fast rate dynamics but unstable for the slow adaptation dynamics, plus it was also susceptible to fluctuation-driven transitions.

### UP and DOWN state statistics in the model

To quantify the relative impact of activity fluctuations and adaptation in causing U-D transitions in the data, we compared the dynamics of the model for different adaptation strengths β and different values of the E effective threshold θ_*E*_. The (θ_*E*_, β) plane was divided into four regions with UD alternations, corresponding to the four combinations of metastability and quasi-stability (Fig. 5A). Since only metastable states tend to give exponentially distributed durations with CV~1, the large variability found in both U and D durations (Fig. 2B) constrained the model to the subregion where both states were metastable and UD and DU transitions were driven by fluctuations (red area in Fig. 5A). The existence of serial correlations between consecutive U and D in the data (Fig. 2C-D) discarded an adaptation-free regime (β = 0), in which transitions were solely driven by fluctuations and the duration of each period was independent of previous durations (Fig. 5B right). Thus, we explored a regime with β > 0 but still in the region where both states were metastable (Fig. 5B, green square) producing alternation dynamics (Fig. 5C top) with broad U and D duration distributions and relatively high CVs (Fig. 5D top). Moreover, the rates showed an autocorrelation function qualitatively similar to the data, with negative side-lobes but no clear traces of rhythmicity (Fig. 5E). Adaptation introduced correlations across consecutive periods (Fig. 5D bottom) because at the transition times the system kept a memory of the previous period in the adaptation value *a(t)*. For adaptation to introduce substantial correlations, *a(t)* had to be variable at the transition times (Lim and Rinzel, 2010), a condition that required adaptation to be fast, to vary within one period, but not too fast to prevent reaching the equilibrium (Fig. 5C bottom trace). Thus, when a strong fluctuation caused a premature UD transition, i.e. a short *U*_*k*_, adaptation had no time to build up and tended to be small, increasing the probability of a premature DU transition in the following D period, i.e. a short *D*_*k*+1_. Conversely, a long *U*_*k*_ recruited strong adaptation that required a long *D*_*k*+1_ to recover (see highlighted examples in Fig. 5C). In this regime, changes in *a(t)* alone did not cause transitions but did modulate the probability that an external fluctuation would cause a transition (Moreno-Bote et al., 2007). Altogether, this analysis suggests that the observed U-D dynamics occurred in a regime with strong random fluctuations, that these fluctuations were necessary to cause the transitions, and that adaptation modulated the timing of the transitions and consequently introduced correlations between the duration of consecutive periods.

**Figure 5.**
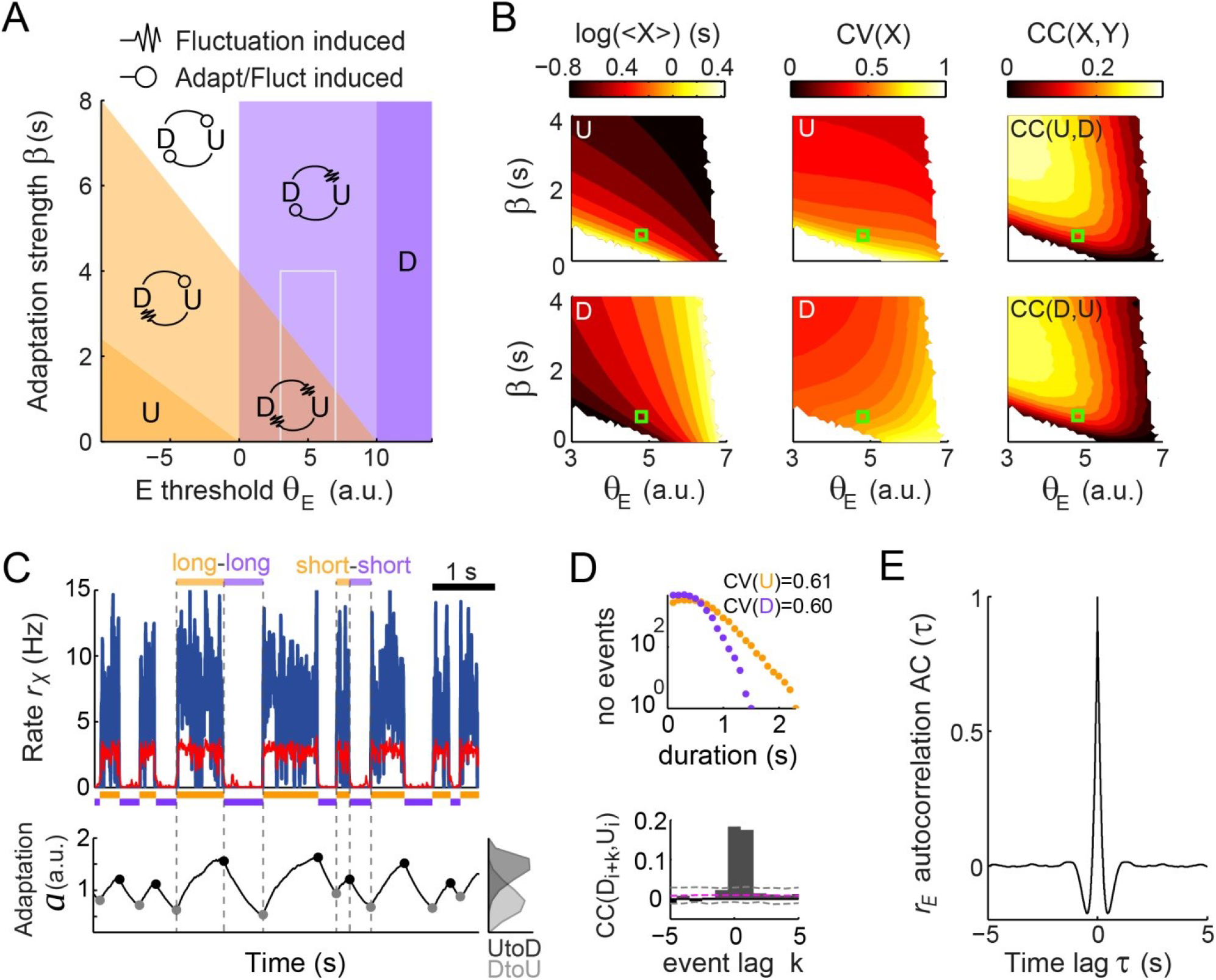
*Fluctuations and weak adaptation are required in the model to explain the U-D statistics of the data.* **(A)** Different dynamical regimes of the model as a function of the adaptation strength β and the effective threshold θ_*e*_. Each U and D state is either meta-stable or quasi-stable depending on whether the transitions to the opposite state can be caused by fluctuations or adaptation+fluctuations, respectively (see arrow code in top inset). There are five region types: regions with a single stable state and no transitions (dark violet and dark orange), a region with both U and D meta-stable (light red), one with both U and D quasi-stable (white) and mixed regions with a meta-stable and a quasi-stable state (light orange and light violet). **(B)** Statistics of U (top) and D (bottom) periods obtained from numerical simulations: mean durations (left), duration CV (center) and of cross-correlation CC of consecutive periods (right) as a function of β and θ_*e*_. The region analyzed is marked in A (gray rectangle). Fluctuations were σ = 3.5. White areas indicate very low transition rate. **(C-E)** Model example quantitatively reproducing some U-D statistics of the data. The β and θ_*e*_ used are marked in B (green square; θ_*e*_ = 4.8 a.u., β = 0.7 Hz^−1^). Example traces of *r*_*e*_*(t)*, *r*_*l*_*(t)*, and *a(t)* show U-D transitions with irregular durations (C). Black and gray filled dots indicate the adaptation values at the UD and DU transition times, respectively. The corresponding histograms illustrate the variability of these values (C bottom right). **(D)** Top: Distributions of U and D period durations. Bottom: Cross-correlograms of D and U periods for different lag values (compare with Fig. 2C). Grey dashed lines show global error bands and magenta dashed line shows mean CC of shuffles. **(E)** Autocorrelogram of *r*_*E*_(t) shows no traces of rhythmicity.

### Dynamics of E and I populations during UP and DOWN states: model and data

According to the model, adaptation currents in the E population can parsimoniously account for the U-D serial correlations but this is in apparent contradiction with the fact that the population rate *R(t)* in the data did not decrease significantly during U periods (Fig. 3C-E). To reconcile these two seemingly contradictory observations we used the model with the parameters that matched the data’s U and D statistics (Fig. 5C-E) to characterize the time course of the rates *r*_*E*_*(t)* and *r*_*I*_*(t)* averaged across DU and UD transitions. Interestingly, the average *r*_*E*_(t) at the beginning and at the end of U periods did not show much difference whereas the average *r*_*l*_(*t*) showed a larger decrease over the U period (Fig. 6A). Thus, although only the E and not the I population included intrinsic adaptation mechanisms, it was *r*_*I*_(*t*) the one that exhibited the most pronounced decay during U periods. This was a direct consequence of the specific conditions that gave rise to bistability in our model: the difference in thresholds, i.e. θ_*l*_ > θ_*E*_, and the fact that the I-nullcline has a higher slope than the E-nullcline (Eq. 21 in Methods). These features imposed that as adaptation built up during U periods, the downward displacement of the E-nullcline caused a greater decrease in *r*_*l*_(*t*) than in *r*_*E*_(*t*) (compare “decay” colored bands in Fig. 6B). With this arrangement the drop in *r*_*E*_(*t*) could be made arbitrarily small by increasing the slope of the I-nullcline (Fig. 6B). During D periods the average *r*_*E*_(*t*) did show a substantial increase due to the recovery of adaptation, whereas the *r*_*l*_(*t*) did not. This was because in the D state, the quiescent network behaved as isolated neurons reflecting the dynamics of intrinsic adaptation which was only present on E cells. In sum, if the majority of the neurons that we recorded experimentally were excitatory, the model could explain why adaptation currents did not cause a significant decrease in the average rate during U periods (Fig. 3C-D). The model in addition predicts that the rate of inhibitory neurons should exhibit a noticeable decrease during U periods.

**Figure 6.**
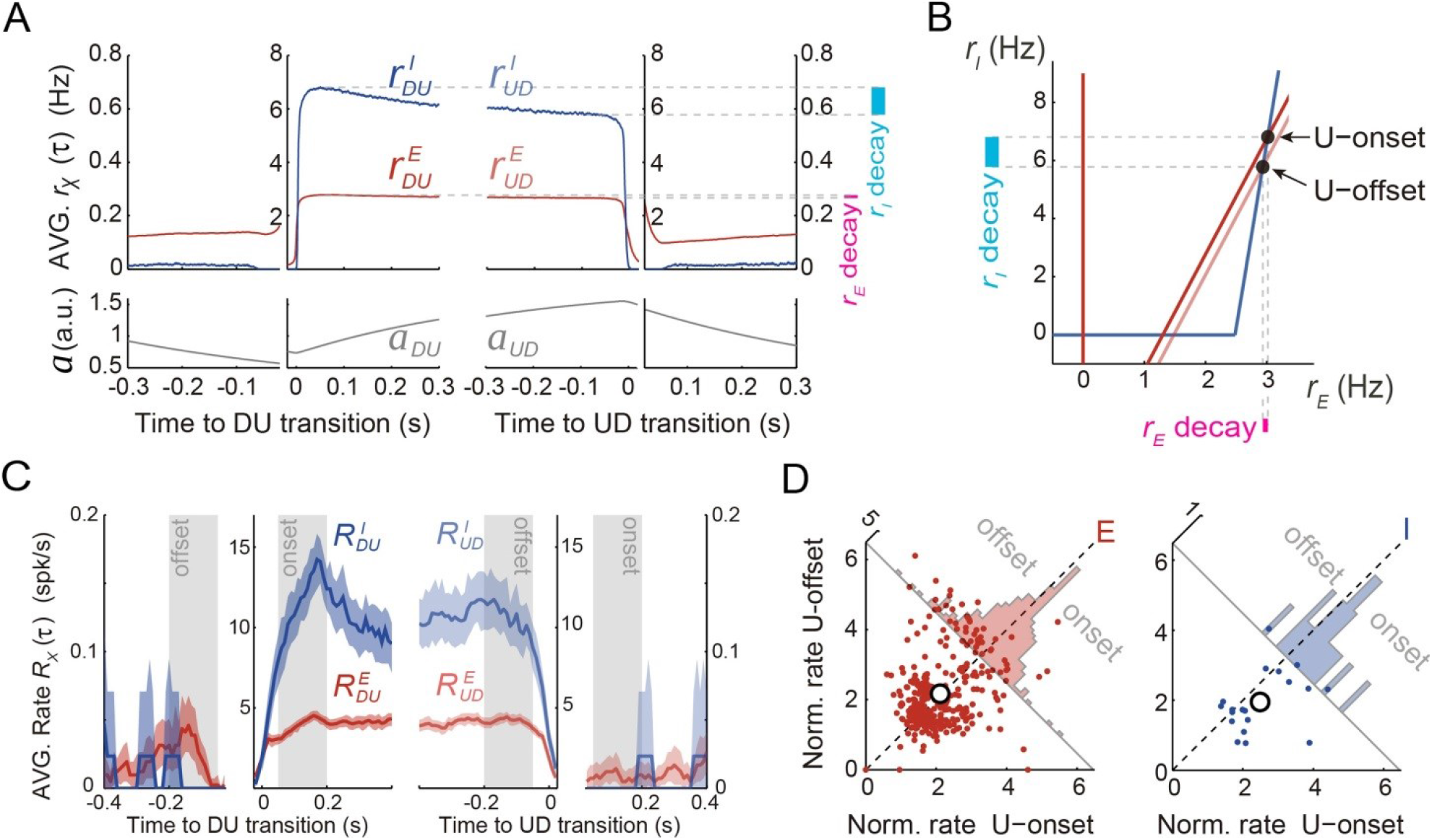
*Excitatory and inhibitory populations during UP and DOWN alternation dynamics.* **(A)** Model average population rates *r*_*e*_and *r*_*l*_ and adaptation *a* as a function of time, aligned at DU and UD transitions (same simulation parameters as in Fig 5C). **(B)** Model predicts a pronounced decay for *r*_*i*_(cyan bar) with minimal decay of *r*_*e*_(pink bar) throughout UP periods, despite adaptation is exclusively included in E cells (Fig 4A). **(C)** Example experiment averaged putative excitatory and inhibitory population rates *(R*^*e*^(t) and *R*^*l*^(t), respectively) aligned at DU and UD transitions. **(D)** Normalized firing rates from individual neurons (see methods) pooled from different experiments (n=5; 330 putative E cells and 21 putative I cells active during U) comparing the activity from putative E and I cells during U onset and offset periods (gray shaded areas from panel C), reveals a significant decrease of I cells during U periods.

Motivated by this prediction, we investigated the dynamics of the rates of excitatory and inhibitory neurons during U and D periods in the experimental data. Based on spike waveforms, isolated units from n=5 experiments were classified into putative interneurons (I) and putative excitatory neurons (E), following previously described procedures (Barthó et al., 2004). The average rate for E and I populations (*R*_*E*_*(t)* and *R*_*I*_*(t)*, respectively) displayed similar profiles across UD alternations, although higher values were observed for I cells during Us (see example experiment in Fig. 6C). To assess the modulation of the rates during U periods, we looked at the normalized individual rates of all the E and I neurons (n=330 and 21, respectively). As predicted by the model (Fig. 6B), I cells displayed a significant rate decay during U periods that was not observed in E cells (Fig. 6E; mixed-effects ANOVA with factors neuron type (E/I), onset/offset and neuron identity and experiment as random factors: interaction neuron type **x** onset/offset *F*(1,349)=6.3, *p*=0.013). During D periods, E cells also showed a significant increase in rate (Wilcoxon signed rank test *P*=0.0092), just like that observed in the whole cell population, whereas no rate change was found in I cells (not shown). Although these changes observed during D periods were also predicted by the model, properly testing the significance of this interaction would require a larger data set with more I cells. The validation of the prediction on the counter-intuitive emergent dynamics of E and I rates during U periods strongly suggests that the mechanism dissected by the model drives the spontaneous dynamics of the cortical circuitry under urethane anesthesia.

## Discussion

Using cortical population recordings we have shown that UP and DOWN period durations are irregular, show positive serial correlation, but there is no significant decrease of population rate during UP periods. These findings seem inconsistent with each other, as some support and others challenge the idea that UP-DOWN dynamics are caused by cell or synaptic adaptive mechanisms. Using a standard EI rate model network, we have proposed a novel bistable regime based only on the expansive threshold non-linearity of the transfer function and on a reported asymmetry between E and I spiking thresholds. While fluctuations produce transitions between the quiescent state (D) and the inhibition-stabilized state of arbitrarily low rate (U), adaptation acting on the E population only facilitates the effect of fluctuations causing the transitions. Paradoxically, because of the asymmetry between E and I thresholds, adaptation causes a marginal decay of E rates but a significant decay of I rates during UP periods. This counterintuitive prediction, specific of our model, was validated in the experimental data.

Adaptive processes constitute the mechanistic hallmark for cortical UP and DOWN dynamics generation (Contreras et al., 1996; Sanchez-Vives and McCormick, 2000; Timofeev et al., 2000). This principle has been used in several computational models, by implementing synaptic short-term depression (Bazhenov et al., 2002; Benita et al., 2012; Hill and Tononi, 2005; Holcman and Tsodyks, 2006; Mejias et al., 2010), or activity-dependent adaptation currents (Compte et al., 2003b; Destexhe, 2009; Latham et al., 2000; Mattia and Sanchez-Vives, 2012).

Consistent with an adaptive process generating the dynamics, UP and DOWN states observed *in vitro* display clear rhythmicity with Gaussian shaped UP and DOWN duration distributions (Mattia and Sanchez-Vives, 2012). An *in vivo* study using ketamine anesthesia in mice reported a reduced UP and DOWN duration variability across multiple cortical areas with CVs around 0.2-0.4 (Ruiz-Mejias et al., 2011). Moreover, a comparison of the UP and DOWN dynamics in the cat observed under ketamine anesthesia and those found in slow wave sleep (SWS) showed that the alternations were more rhythmic under ketamine (Chauvette et al., 2011). In contrast, our data displayed large variability (mean CV(U)~CV(D)≃0.7) and skewed distributions of UP and DOWN period durations (Fig. 2B), in agreement with previous studies using urethane anesthesia (Dao Duc et al., 2015; Stern et al., 1997). Although a direct comparison between the UP-DOWN dynamics under urethane anesthesia and during natural sleep has not been made, urethane seems to mimic sleep in several aspects. First, it induces spontaneous alternations between synchronized and desynchronized states (Curto et al., 2009; Steriade et al., 1994), resembling the alternations between SWS and REM sleep (Clement et al., 2008; Whitten et al., 2009). Second, the irregular UP-DOWN transitions observed under urethane anesthesia resemble the variability observed in SWS (Ji and Wilson, 2007; Johnson et al., 2010). Preliminary analysis of rat and mouse prefrontal cortex during SWS with the same population-based U-D detection methods used here (Methods) showed that U periods had comparable mean length but were more irregular (CV~1) than under urethane anesthesia (Fig. 1B) whereas D periods were shorter (mean ~150 ms) and slightly more regular (CV~0.5) (unpublished observations). Such an asymmetry in the duration and irregularity of U-D periods can be easily reproduced in our model by choosing parameters in the mixed region where U is meta-stable and D is quasi-stable (Fig. 5A light orange). In addition, we found non-zero correlations between consecutive D-U and U-D period durations, a feature that was not observed previously in similar experimental conditions (Stern et al., 1997). Reduced statistical power (~30 U-D/D-U pairs were considered by (Stern et al., 1997) versus a range of 462-758 pairs in our n=7 experiments) and different U-D detection methods (intracellular membrane potential thresholding) could be the reasons for this discrepancy.

### Bistability in cortical networks at low firing rates

Bistability in a dynamical system refers to the coexistence of two possible steady states between which the system can alternate (Angeli et al., 2004). This principle has been used to interpret UP and DOWN states as two attractors of cortical circuits (Cossart et al., 2003; Shu et al., 2003) and it seems to underlie higher cognitive functions (Compte, 2006; Durstewitz, 2009). In particular, multi-stability in recurrent cortical networks has been postulated to underlie the persistent activity observed during the delay period in working memory tasks. Extensive theoretical work has shown that based on the change in curvature of the neuronal *f -* I curve, i.e. from expansive to contractive, recurrent network models generate two types of co-existing attractors: a spontaneous state with arbitrarily low rates (falling in the expansive part of the *f* - I curve) and a sustained activity attractor where the reverberant activity of a subpopulation of neurons could be maintained at a rate on the contractive part of the *f* - I curve (Amit and Brunel, 1997; Brunel, 2000a; Wang, 2001). Thus, unless additional mechanisms are included, e.g. synaptic short-term depression and facilitation (Barbieri and Brunel, 2007; Hansel and Mato, 2013; Mongillo et al., 2012) or fined-tuned EI balance (Renart et al., 2007), the rate of persistent states is lower-bounded by the rate where the *f -* I curve changes from convex to concave (~10-20 spikes/s). Moreover, because of this the sustained attractor operates in an unbalanced supra-threshold regime where spike trains tend to be more regular (i.e. lower inter-spike-interval CV, (Barbieri and Brunel, 2007; Hansel and Mato, 2013; Renart et al., 2007)) than those observed in the data (Compte et al., 2003b).

UP and DOWN states represent in contrast transitions between very different levels of activity: a quiescent state and a very low rate state. Given that we recorded neurons extracellularly, our estimate of the mean firing rate during UP periods (3.7 spikes/s) is most likely an overestimation. Whole cell intracellular recordings have reported rates in the range 1-2 spikes/s (Constantinople and Bruno, 2011), 0.4 spikes/s in Pyramidal L2/3 of the somatosensory cortex of awake mice (Gentet et al., 2012), 0.1 spikes/s Pyramidal L2/3 cells in somatosensory cortex during UP periods in anesthetized rats (Waters and Helmchen, 2006), or 0.1-0.3 spikes/s in V1 of awake mice (Haider et al., 2013). Juxtacellular recordings have found values near 4-5 spikes/s (Massi et al., 2012; Sakata and Harris, 2009) whereas Calcium imaging experiments report spontaneous rates <0.1 spikes/s (Kerr et al 2005). Despite UP rates being so low, rate models have commonly used the change in curvature of the transfer function to generate UP and DOWN dynamics (Curto et al., 2009; Lim and Rinzel, 2010; Mattia and Sanchez-Vives, 2012; Mochol et al., 2015). It is also for this reason that most spiking network models generating UP and DOWN transitions exhibit unrealistically high rates during U periods (in the range 10-40 spikes/s) with relatively regular firing (Bazhenov et al., 2002; Compte et al., 2003b; Destexhe, 2009; Hill and Tononi, 2005).

An alternative mechanism to generate bistability between UP and DOWN states has been the shunting or divisive effect of inhibitory synaptic conductances, a mechanism that can produce nonmonotonic transfer functions and yield bistability between a zero rate state and a state of very low rate (Kumar et al., 2008; Latham et al., 2000; Vogels and Abbott, 2005). Latham and colleagues (Latham et al., 2000) addressed the question of how to obtain a state of low firing rates (i.e. <1 spikes/s) in a recurrent EI network and concluded that there were two alternative mechanisms: the most robust was to have a single attractor that relied on the excitatory drive from endogenously active neurons in the network or from external inputs. In fact, excitatory external inputs have been widely used to model low rate tonic spontaneous activity (i.e. no DOWN states) in EI networks of current-based spiking units (Amit and Brunel, 1997; Brunel, 2000b; Vogels and Abbott, 2005). Alternatively, in the absence of endogenous or external drive, a silent attractor appears and a second attractor can emerge at a low rate over a limited range of parameters if inhibition exerts a strong divisive influence on the excitatory transfer function (Latham et al., 2000). Based on this, a spiking network of conductance-based point neurons with no external/endogeneous activity could alternate between UP (0.2 spikes/s) and DOWN (0 spikes/s) periods via spike frequency adaptation currents.

Our model proposes a more parsimonious mechanism underlying UP-DOWN bistability: the ubiquitous expansive threshold non-linearity of the transfer function plus the asymmetry in threshold (θ_*I*_ > θ_*E*_) and gain (larger for I than E cells). We used a threshold-linear function for simplicity but other more realistic choices (e.g. threshold-quadratic) produced the same qualitative results. The threshold asymmetry is supported by *in vitro* patch clamp experiments revealing that firing threshold of inhibitory fast-spiking neurons, measured as the lowest injected current causing spike firing, is higher than that of excitatory regular-spiking neurons (Cruikshank et al., 2007; Schiff and Reyes, 2012). Thus, inhibition in this model becomes active when external inputs onto E cells during the DOWN state are strong enough to push the system above the separatrix and ignite the UP state. Once recruited, inhibition is necessary to stabilize the activity because, in its absence, the positive feedback would make the UP state unstable. These are the conditions that define an Inhibitory Stabilized Network (Ozeki et al., 2009), which in large networks is referred to as the balance state (Amit and Brunel, 1997; Renart et al., 2010; van Vreeswijk and Sompolinsky, 1998). Thus, according to our model, cortical circuits alternate between a quiescent state with no activity and a state of balanced irregular and asynchronous low rate activity (Renart et al., 2010). It is for this reason that in the rodent, mean pairwise spiking correlations during UP periods are negligible (Renart et al., 2010; Stern et al., 1998) and that the rate of coordinated transitions to the DOWN state predicts the magnitude of correlations across different brain states (Mochol et al., 2015).

A direct implication of the bistability obtained in our model was that intrinsic adaptation of excitatory neurons (McCormick et al., 1985) did not cause a noticeable decrease in *r*_*E*_ during the UP periods but instead produced a significant decay in the inhibitory rate *r*_*l*_. We confirmed this prediction in our data (Fig. 6C-D). Interestingly, the same effect was also observed in ketamine anesthetized animals from both extracellular (Luczak and Barthó, 2012) and intracellular recordings resolving synaptic conductances (Haider et al., 2006). During DOWN periods in contrast, the network is not in a balanced state and recovery from adaptation caused a significant increase in the rate of putative excitatory neurons, as predicted by the model. In sum, our results present the first EI network model with linearly summed inputs exhibiting bistability between a quiescent state and a balanced state with arbitrary low rate.

### The role and origin of fluctuations in UP-DOWN switching

Our findings stress the role of input fluctuations inducing transitions between the UP and DOWN network attractors because noise-induced alternations generate periods with large variability as found in the data (Fig. 2A-B). Adaptation was also necessary to introduce positive serial correlations and to reproduce the observed gamma-like UP-DOWN distributions (compare Fig. 2A-B and Supp. Fig 1 with Fig. 5D) because it caused a soft refractory period after each transition decreasing the duration CVs below one (Fig. 2B). In our network model fluctuations were simply introduced by a time-varying Gaussian input so that in both DU and UD transitions the noise had the same external origin. In cortical circuits however these two transitions are very different: while in UP-DOWN transitions the fluctuations can originate in the stochasticity of the spiking activity during the UP period, DU transitions depend on either local circuit mechanisms that do not need spiking activity or on external inputs to escape from a quiescent state. Previous bistable spiking network models have used the stochasticity of the recurrent spiking activity to cause transitions from a balanced regime into a quiescent state (i.e. UP-to-DOWN) (Kumar et al., 2008; Latham et al., 2000). Other models have proposed that synaptic noise (e.g. spontaneous miniatures) could cause the transitions from the quiescent state (i.e. DOWN to UP) (Bazhenov et al., 2002; Holcman and Tsodyks, 2006; Mejias et al., 2010). Preliminary analysis using a spiking EI network to produce UP-DOWN alternations shows that to cause noise-driven transitions from a quiescent state synaptic inputs into each neuron need to be correlated and non-Gaussian (not shown). Gaussian uncorrelated inputs must generate an unrealistically high DOWN firing rate in order to yield a measurable escape probability to the UP state (because escape requires the synchronous occurrence of multiple independent neuronal discharges). That is why synaptic noise, which is in principle independent across synapses, could not account for DOWN-to-UP transitions having variable DOWN duration and near zero firing, e.g. CV(D)~0.7 (Fig. 2B) and *r*_*D*_ ~ 0.018 spikes/s (Supp. Fig. 2B). This reasoning favors instead synchronous external input *kicks* as the inducers of DOWN-to-UP transitions. Evidence for such temporally sparse synchronous inputs comes from intracellular membrane potential recordings under some types of anesthesia (penthobarbital or halothane) showing «presynaptic inputs […] organized into quiescent periods punctuated by brief highly synchronous volleys, or “bumps”» (DeWeese and Zador, 2006). We postulate that these spontaneous bumps (DeWeese and Zador, 2006; Tan et al., 2013; Taub et al., 2013) (1) are caused by synchronous external inputs impinging on the neocortex, possibly from thalamocortical neurons (Crunelli and Hughes, 2010), since spontaneous bumps resemble sensory evoked responses (DeWeese and Zador, 2006) or from hippocampal Sharp Wave Ripples (Battaglia et al., 2004); (2) their timing resembles a Poisson stochastic process rather than a rhythmic input (Tan et al., 2013); (3) they lie at the origin of the DOWN-to-UP transitions that we observe. Despite the fact that UP-DOWNlike activity can emerge in cortical slices *in vitro* (Cossart et al., 2003; Fanselow and Connors, 2010; Mann et al., 2009; Sanchez-Vives and McCormick, 2000) the intact brain can generate more complex UP-DOWN patterns than the isolated cortex, with subcortical activity in many areas correlating with transition times (Battaglia et al., 2004; Crunelli et al., 2015; Crunelli and Hughes, 2010; David et al., 2013; Lewis et al., 2015; Slézia et al., 2011; Ushimaru et al., 2012). Further analysis using detailed spiking models will be necessary to characterize the detailed conditions under which external inputs can trigger an UP state, such as the number of cortical spikes that must be evoked and their degree of synchrony, the role of inhomogeneities in the connectivity generating trigger “hot spots” (Tsodyks et al., 2000) and stereotyped onset patterns (Luczak et al., 2009; Roxin et al., 2008).

These arguments bring forward the idea that DOWN to UP transitions are, at least in part, caused by punctuated external synchronous inputs (Battaglia et al., 2004; Johnson et al., 2010), with slow intrinsic adaptation mechanisms contributing to modulate the probability that these events trigger a transition (Moreno-Bote et al., 2007). This complements the idea that UP-DOWN dynamics reflect an endogenous oscillation of the neocortex and connects to the role of UP-DOWN states in memory consolidation: because in the active attractor (UP) the *stationary* activity is irregular and asynchronous (Renart et al 2010), the existence of a silent attractor enables synchronous transient dynamics in the form of DOWN to UP transitions. These transients generate precise temporal relations among neurons in a cortical circuit (Luczak et al 2007), which can cause synaptic plasticity underlying learning and memory (Peyrache et al 2009). We speculate that, while the transient dynamics are triggered by external inputs, adaptation, by introducing refractoriness in this process, parses transition events preventing the temporal overlap of information packets (Luczak et al., 2015).

## Supplementary figures

**Supplementary Figure 1.**
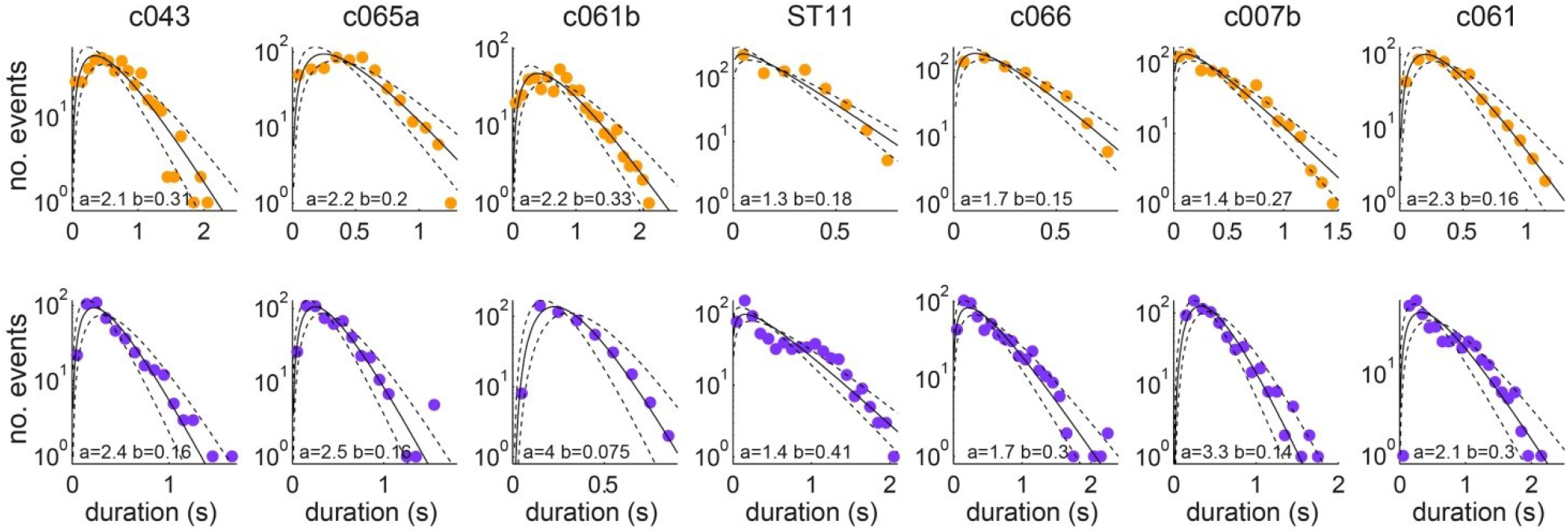
Distributions of U (top row) and D (bottom row) period durations for the seven experiments (columns). Gamma fit for each distribution is shown in black. The shape *a* and scale *b* parameters of the Gamma fit are displayed at the bottom of each panel. 95% CI of the fit displayed in dashed line.

**Supplementary Figure 2.**
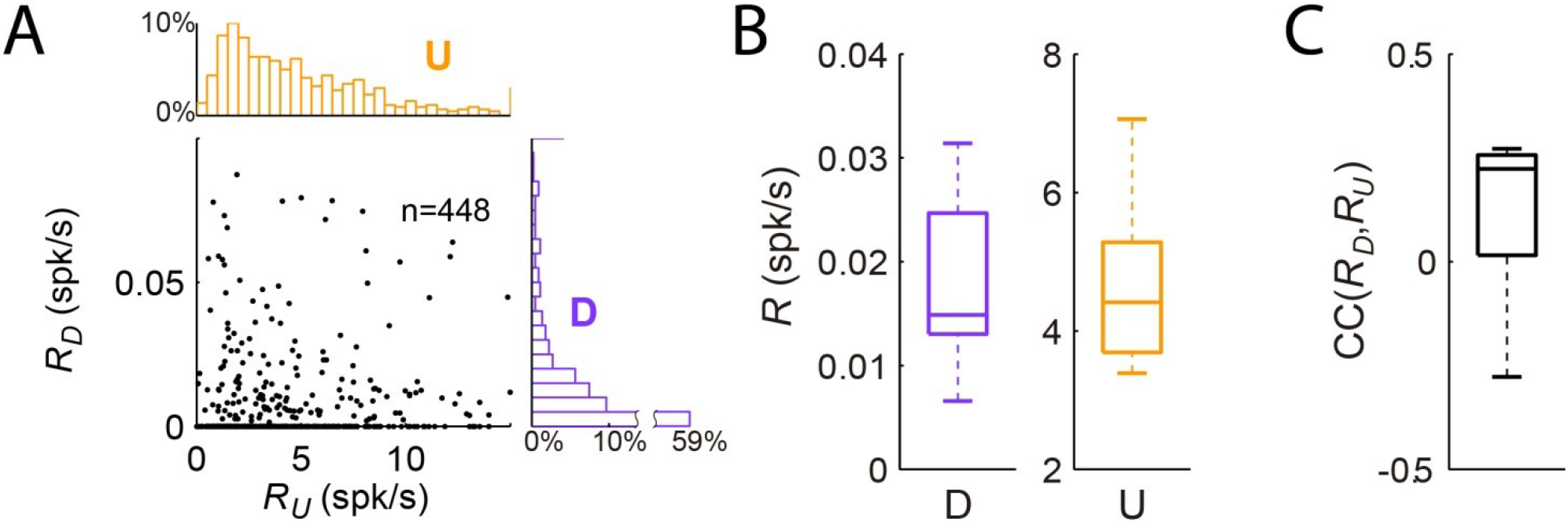
*Spiking statistics of individual units during U and D periods.* **(A)** Scatter plot of average firing rate during U periods versus firing rate during D periods (R_U_ vs R_D_), from all the isolated cells from the different experiments (n=7 animals; 448 cells). Marginal distribution of firing rates for R_U_ in orange and R_D_ in violet. **(B)** Statistics of mean single cell firing rates during D (left boxplot) and U (right boxplot) periods for different experiments. **(C)** Statistics of mean correlation between firing rate during D and U periods for different experiments.

## Materials and methods

### Experimental procedures

Adult, male Sprague-Dawley rats (250-400g) were anesthetized with urethane (1.5 g/kg) and supplemental doses of 0.15 g/kg were given when necessary after several hours since the initial dose. We also used an initial dose of Ketamine (15-25 mg/kg) before the surgery to induce the anesthetized state quickly. We then performed a craniotomy over the somatosensory cortex, whose position was determined using stereotaxic coordinates. Next 32 or 64 channels silicon microelectrodes (Neuronexus technologies, Ann Arbor MI) were slowly inserted into in deep layers of the cortex (depth 600-1200 μm; lowering speed ~ 1mm/hour). Probes had either eight shanks each with eight staggered recording sites per shank (model Buzsaki64-A64), or four shanks with two tetrode configurations in each (model A4×2-tet-5mm-150-200-312-A32). Neuronal signals were high-pass filtered (1Hz) and amplified (1,000X) using a 64-channel amplifier (Sensorium Inc., Charlotte, VT), recorded at 20kHz sampling rate with 16-bit resolution using a PC-based data acquisition system (United Electronic Industries, Canton, MA) and custom written software (Matlab Data Acquisition Toolbox, MathWorks) and stored on disk for further analysis.

### Data Analysis

Spike sorting was performed using previously described methods (Harris et al., 2000). Briefly, units were isolated by a semiautomatic algorithm (http://klustakwik.sourceforge.net) followed by manual clustering procedures (http://klusters.sourceforge.net). We defined the *Population activity* as the merge of the spike trains from all the well isolated units.

#### Putative E/I neuronal classification

Isolated units were classified into narrow-spiking (I) and broad-spiking (E) cells based on three features extracted from their mean spike waveforms: spike width, asymmetry and through-to-peak distance. The two classes were grouped in the space of features by k-means clustering (Barthó et al., 2004; Csicsvari et al., 1998; Sirota et al., 2008).

#### Synchronized state assessment

We classified the brain state based on the silence density defined as the fraction of 20 ms bins with zero spikes in the Population activity in 10 s windows (Mochol et al., 2015; Renart et al., 2010). Epochs with consecutive windows of silence density above 0.4, standard deviation below 0.1 and longer than 5 min, were considered as sustained synchronized brain state and were used for further analysis (synchronized states durations mean±SD: 494±58 s, n=7 epochs).

#### UP & DOWN transitions detection

UP-DOWN phases have been commonly defined from intracellular recordings by detecting the crossing times of a heuristic threshold set on the membrane potential of individual neurons (Mukovski et al., 2007; Stern et al., 1997), or from local field potential signals (Compte et al., 2008; Mukovski et al., 2007) or combined together with the information provided by multi-unit activity (Haider et al., 2006; Hasenstaub et al., 2007). Defining UP-DOWN phases from single-unit recordings is more challenging because individual neurons fire at low rates discharging very few action potentials on each UP phase (Constantinople and Bruno, 2011; Gentet et al., 2012; Waters and Helmchen, 2006). However, pooling the spiking activity of many neurons into a population spike train reveals the presence of co-fluctuations in the firing activity of the individual neurons and allows accurate detection of UP-DOWN phases (Luczak et al., 2007; Saleem et al., 2010). We used a discrete-time hidden semi-Markov probabilistic model (HMM) to infer the discrete two-state process that most likely generated the population activity (Chen et al., 2009). Thus, the population activity spike count was considered as a single stochastic point processes whose rate was modulated by the discrete hidden state and the firing history of the ensemble of neurons recorded. In order to estimate the hidden state at each time, the method used the expectation maximization (EM) algorithm for the estimation of the parameters from the statistical model (Chen et al., 2009). Although the discrete-time HMM provides a reasonable state estimate with a rather fast computing speed, the method is restricted to locate the UP and DOWN transition with a time resolution given by the bin size (*T*) for the population activity spike count (10 ms in our case). The initial parameters used for the detection were: Bin-size *T* = 10 ms, number of history bins J=2 (sets the length of the memory, i.e. J=0 is a pure Markov process); history-dependence weight β = 0.01 (i.e. β=0 for a pure Markov process); transition matrix P_DU_=P_UD_=0.9, P_DD_=P_UU_=0.1; rate during UP periods α = 3, and rate difference during DOWN and UP periods μ = -2 (Chen et al., 2009). The algorithm gives an estimate of the state of the network on each bin *T*. Adjacent bins in the same state are then merged to obtain the series of *putative UP (U) and DOWN (D)* periods. The series is defined by the onset 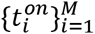 and offset 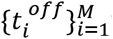 times of the Us, where M is the total number of Us, that determine the *i*-th UP and DOWN period durations as (see Fig. 1C):

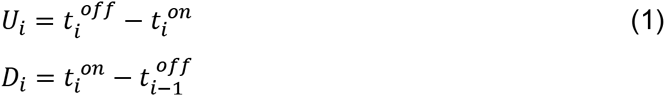

#### Statistics of UP & DOWN durations

The mean and the coefficient of variation of *U*_*i*_ were defined as

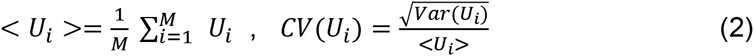

where:

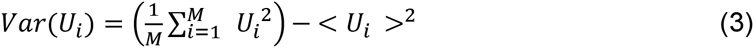

and equivalently for <*D*_*i*_> and *CV*(*D*_*i*_). We controlled whether variability in *U*_*i*_ was produced by slow drifts by computing *CV*_*2*_ a measure of variability not contaminated by non-stationarities of the data (Compte et al., 2003a; Holt et al., 1996).

The serial correlation between *U*_*i*_ and *D*_*i*+*k*_, with *k* setting the lag in the U-D series, e.g. *k* =0 (*k* = 1) refers to the immediately previous (consecutive) DOWN period, was quantified with the Pearson correlation coefficient defined as:

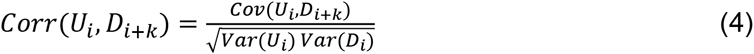

where the covariance was defined as:

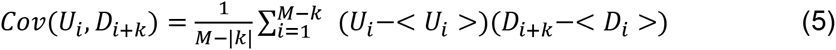

Values of *U*_*i*_ and *D*_*i*_differing more than 3 standard deviations from the mean were discarded from the correlation analysis (circles in Fig 2C). To remove correlations between *U*_*i*_ and *D*_*i*_produced by slow drifts in the durations we used resampling methods developed to remove slow correlations among spike trains (Amarasingham et al., 2012). We generated the *l-th* shuffled series of U periods 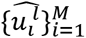 by randomly shuffling the order of the Us in the original series 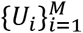 within intervals of 30 s. The same was done to define the shuffled series of D periods 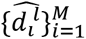. The two shuffled series lack any correlation except that introduced by co-variations in the statistics with a time-scale slower than 30 s. We generated *L*=1000 independent shuffled series 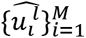 and 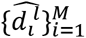 with *l*=1,2,…*L*, computed the covariance 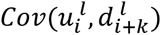 for each and the averaged over the ensemble 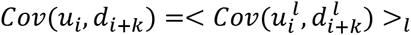. Finally, the correlation due to co-fluctuations of Us and Ds faster than 30 s was computed by subtracting *Cov*(*u*_*i*_, *d*_*i*+*k*_) from*Cov*(*U*_*i*_, *D*_*i*+*k*_) in Eq. 5. Significance of the correlation function *Corr*(*U*_*i*_, *D*_*i*+*k*_) was assessed by computing a point-wise confidence interval from a distribution of *L* correlograms 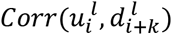, for *I*=1…*L* (*L*=10000), computed from each shuffled series the same way as for the original series (gray dashed bands in Fig 2C). To take into account multiple comparisons introduced by the range in lag *k*, we obtained *global* confidence intervals (black dashed bands in Fig 2C) by finding the *P* of the pointwise intervals for which only a fraction *0.05* of the correlograms 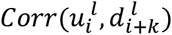 crosses the interval bands at *any* lag *k*= −7…7 (see (Fujisawa et al., 2008) for details).

#### Spike count statistics

We divided the time in bins of *dt*=1 ms and defined the spike train of the *j*-th neuron as:

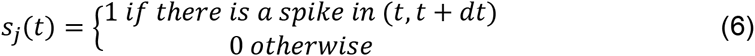

The *spike count* of the *j*-th neuron over the time window (*t*-T/2, *t*+T/2) was obtained from

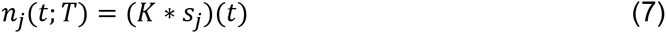

where * refers to a discrete convolution and *K*(*t*) is a square kernel which equals one in (−T/2,T/2) and zero otherwise.

The instantaneous rate of the *j*-th neuron was defined as:

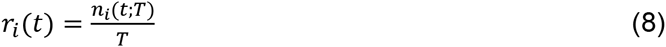

and therefore the instantaneous population rate was defined as:

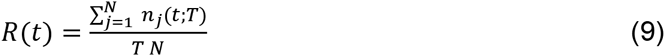

where N is the total number of well isolated and simultaneously recorded neurons. We have dropped the dependence on *T* from *r*_*i*_(*t*) and *R*(*t*) to ease the notation. We also defined the instantaneous E-population and I-populations rates, *R*^*E*^(*t*) and *R*^*I*^(*t*) respectively, as those computed using cells in the E and I subpopulations separately.

#### Population firing statistics during Us and Ds

The instantaneous population rate averaged across Us and Ds and aligned at the D to U transition (DU) was defined as:

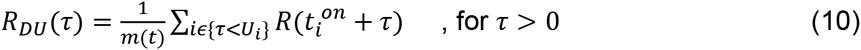

where *τ* is the time to the DU transition. Because Us had different durations, for each *τ* > 0, the sum only included the onset time 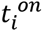 if the subsequent period was longer than *τ* < *U*_*i*_. By doing this we remove the trivial decay we would observe in *R*_*DU*_(*τ*) as *τ*increases due to the increasing probability to transition into a consecutive period *D*_*i*+1_. For *τ* < 0, *R*_*DU*_(*τ*) reflecting the population averaged rate during the Ds, is obtained as in Eq. 10 but including the times 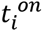 in the sum if the previous D was longer than |*τ*| < *D*_*i*−1_. Similarly, the average population rate aligned at the offset *R*_*UD*_(*τ*) was defined equivalently by replacing 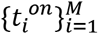 by the series of offset times 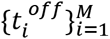. We also defined the onset and offset-aligned averaged population rate for excitatory (E) and inhibitory (I) populations, termed 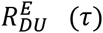 and 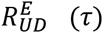 for the E case and similarly for the I case. Moreover, the onset and offset-aligned averaged rate of the *i*-th neuron 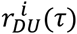 and 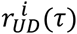 were defined similarly using the individual rate defined in Eq. 8.

The autocorrelogram of the instantaneous population rate was defined as:

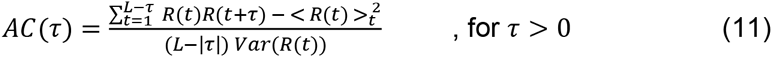

with the sum in *t* running over the *L* time bins in a window of size W. The average < *R*(*t*) >_*t*_ and variance were performed across time in the same window. To avoid averaging out a rhythmic structure in the instantaneous population rate due to slow drift in the oscillation frequency, we computed *AC*(*τ*) in small windows W=20 s thus having a more instantaneous estimate of the temporal structure. With the normalization used, the autocorrelograms give *AC*(*τ* = 0) = 1 and the values with *τ* > 0 can be interpreted as the Pearson correlation between the population rate at time *t* and the population rate at time *t* + *τ* (Fig. 1D).

#### Instantaneous rates at onset and offset intervals

To compare the population rates at the U-onset and U-offset (Fig 3 and 6), we computed for each neuron the mean of 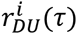 over the window *τ* = (50,200) *s* (U-onset) and the mean of 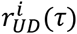 over the window *τ* = (−200, −50) *s* (U-offset). We positioned the windows 50 ms away of the DU and UD transitions in order to preclude the possibility of contamination in the mean rate estimations due to possible misalignments from the U and D period detections. In the averaging we used U and D periods longer than 0.5 s, so that onset and offset windows were always non-overlapping. Equivalent D-onset and D-offset windows were defined in order to compare individual rates during D periods. To make the distribution of mean rates across the cell population Gaussian, we normalized each the rates 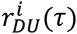 and 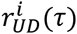 by the overall time-averaged rate of the neuron *r*_*i*_=< *r*_*i*_(*t*) >_*t*_ finally obtaining onset and offset-aligned *normalized* averaged rates (e.g. 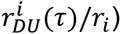. Despite this normalization, the distribution of the normalized rates in the D-onset and D-offset was non-Gaussian (most neurons fired no spikes). Thus we used the non-parametric two-sided Wilcoxon signed rank test to compare onset and offset rates (Fig. 3E). To test the rates changes during U periods in E and I neurons we used a four-way mixed-effects ANOVA with fixed factors onset/offset, E/I and random factors neuron index and animal. We compared the distribution of *normalized* averaged rate difference at the U-onset minus the U -offset (Fig. 3E right, dark gray histogram) with a distribution obtained from the same neurons but randomly shuffling the onset and offset labels of the spike counts but preserving trial and neuron indices (Fig. 3E right, light gray bands show 95% C.I. of the mean histograms across 1000 shuffles). This surrogate data set represents the hypothesis in which none of the neurons shows any onset vs offset modulation. The comparison shows that there are significant fractions of neurons showing a rate decrease and increase that compensate to yield no significant difference on the population averaged rate. The same procedure was followed with the normalized rates in the D-onset and D-offset but the limited number of non-zero spike counts limited the analysis yielding inconclusive results (Fig. 3E left).

### Computational Rate Model

We built a model describing the rate dynamics of an excitatory (*r*_*E*_) and inhibitory population (*r*_*I*_) recurrently connected that received external inputs (Wilson and Cowan, 1972). In addition, the excitatory population had an additive negative feedback term, *a*(*t*), representing the firing adaptation experienced by excitatory cells (McCormick et al., 1985). The model dynamics were given by:

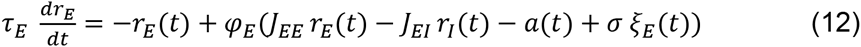

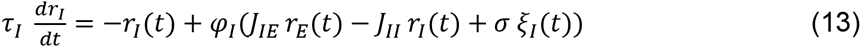

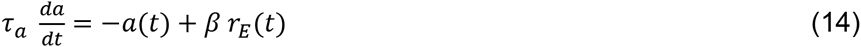

The time constants of the rates were *τ*_*E*_ = 10 ms and *τ*_*I*_ = 2 ms, while the adaptation time constant was *τ*_a_ = 500 ms. The synaptic couplings *J*_*XY*_ > *0* (with *X,Y* = E, I), describing the strength of the connections from *Y* to *X*, were *J*_*EE*_ = 5, *J*_*EI*_ = 1, *J*_*IE*_ = 10, *J*_*II*_ = 0.5 s. Because we are modeling low rates, the adaptation grows linearly with *r*_*E*_ with strength *β*= 0.5 s. The fluctuating part of the external inputs *σ ξ*_*x*_(*t*) was modeled as two independent Ornstein–Uhlenbeck processes with zero mean, standard deviation *σ*= 3.5 and time constant 1 ms for both E and I populations. Because population averaged firing rates during spontaneous activity fell in the range 0-10 spikes/s, we modeled the transfer functions φ_*X*_ as threshold-linear functions:

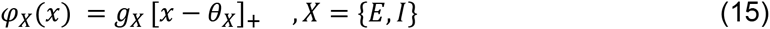

where the square brackets denote [*z*]_+_ = *z* if *z* > 0and zero otherwise, the gains were *g*_*E*_ = 1 Hz and *g*_*I*_ = 4 Hz and the *effective* thresholds *θ*_*E*_ and *θ*_*I*_ represented the difference between the activation threshold minus the mean external current into each population. We took *θ*_*I*_*=* 25 a.u. and explored varying *θ*_*E*_ over a range of positive and negative values (Fig. 5A-B). The choice of thresholds *θ*_*E*_< *θ*_*I*_and gains*g*_*E*_ < *g*_*I*_ reflecting the asymmetry in the *f-I* curve of regular spiking neurons (E) and fast spiking interneurons (I) (Cruikshank et al., 2007; Nowak et al., 2003; Schiff and Reyes, 2012), facilitated that the model operated in a bistable regime (see below).

Input-output transfer functions are typically described as sigmoidal-shaped functions (Haider and McCormick, 2009), capturing the nonlinearities due to spike threshold and firing saturation effects. Since we are interested in modeling spontaneous activity where average population rates are low, we constrained the transfer functions to exhibit only an expanding non-linearity reflecting the threshold and thus avoid other effects that can only occur at higher rates (the contracting non-linearity tends to occur for rates > 30 spikes/s (Anderson et al., 2000; Houweling et al., 2010; Nowak et al., 2003; Priebe and Ferster, 2008). In particular, we modeled φ_X_ as piecewise linear (Schiff and Reyes, 2012; Stafstrom et al., 1984) but the same qualitative bistable regime can be obtained by choosing for instance a threshold-quadratic function. The model equations (Eqs. 12-14) were numerically integrated using a fourth-order Runge-Kutta method with integration time step dt = 0.2 ms. U and D periods in the model were detected by threshold-based method, finding the crossing of the variable *r*_*E*_ with the boundary 1 Hz, where periods shorter than minimum period duration of 50 ms were merged with neighboring periods (small changes in threshold and period durations did not affect qualitatively the results).

#### Fixed points and stability

We start by characterizing the dynamics of the system in the absence of noise. Assuming that the rates evolve much faster than the adaptation, i.e. *τ*_*E*_, *τ*_*I*_ ≪ *τ*_*a*_, one can partition the dynamics of the full system into (1) the dynamics of the rates assuming adaptation is constant, (2) the slow evolution of adaptation assuming the rates are constantly at equilibrium. Thus, the equations of the *nullclines* of the 2D rate dynamics at fixed a, can be obtained from the 2D system given by Eqs. 12-13. The nullclines of this reduced 2D system are obtained by setting its left hand side to zero:

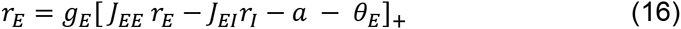

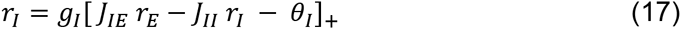

The intersection of the nullclines define the fixed points (*r*\*_*E*_ (*a*), *r*\*_*I*_ (*a*)) of the 2D system to which the rates evolve. Once there adaptation varies slowly assuming that the rates are maintained at (*r*\*_*E*_ (*a*), *r*\*_*I*_ (*a*)) until it reaches the equilibrium at *a* = *βr*\*_*E*_ (*a*).

The network has a fixed point in (*r*_*E*_, *r*_*I*_, *a*)=(0,0,0) if and only if θ_*E*_ *≥* 0 and θ_*I*_ *≥* 0, i.e. when the mean external inputs are lower than the activation thresholds. The stability of this point, corresponding to the DOWN state, further requires θ_*E*_>0, thus preventing the activation of the network due to small (infinitesimal) fluctuations in *r*_*E*_. To find an UP state fixed point with non-zero rates we substitute in Eqs. 16-17 the value of adaptation at equilibrium *a* = *βr*_*E*_, assume the arguments of []_+_ are larger than zero and solve for (*r*_*E*_, *r*_*I*_), obtaining:

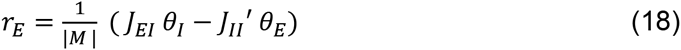

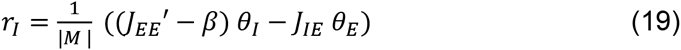

where 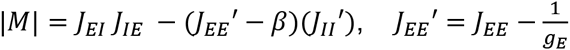 and 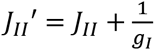.

The conditions for this UP state solution to exist are derived from imposing that the right hand side of Eqs. 18-19 is positive. The stability of this solution (Eq. 21 below) implies that the determinant |*M*| is positive and that if *r*_*I*_ is positive, then *r*_*E*_ is also positive. Thus, provided the stability (Eqs. 21-22), the only condition for the solution to exist is that the right hand side of Eq. 19 is positive:

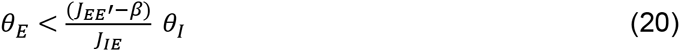

Given the separation of time scales described above, this fixed point is stable if the eigenvalues of the matrix of coefficients of Eqs. 16 and 17 without the term *a* (that we assume is constant) have all negative real part. Because the coefficients matrix is 2 × 2, this is equivalent to impose that the determinant of the matrix has a positive determinant and a negative trace. These conditions yield the following inequalities, respectively:

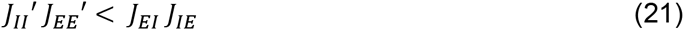

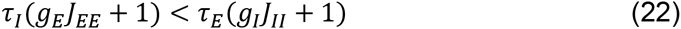

Equation 21 is equivalent to the condition that the I-nullclines of the 2D reduced system has a larger slope than the E-nullcline. From the U existence condition in Eq. 20 and D stability condition, it can also be derived that *J*_*EE*_′ > 0, implying that at fixed inhibition, the E-subnetwork would be unstable (i.e. slope of the E-nullcline is positive). In sum, the conditions for the existence of two stable U and D states imply that the U state would be unstable in the absence of feedback inhibition but the strength of feedback inhibition is sufficient to stabilize it. These are precisely the conditions that define an Inhibitory Stabilized Network state (Ozeki et al., 2009).

#### Phase plane analysis

In this section we determine the different operational regimes of the network in the (*θ*_*E*_, β)-plane (Fig. 5A). In the absence of noise, given that θ_*I*_ ≥ 0, a stable D state exists in the semi-plane (Fig. 5A, violet and red regions):

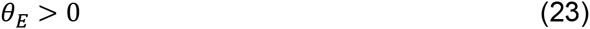

Provided that our choice of synaptic couplings *J*_*xγ*_ and time constants hold the stability conditions (Eqs. 21-22), the U state is stable in the semi-plane given by Eq. 20 (Fig. 5A, orange and red regions):

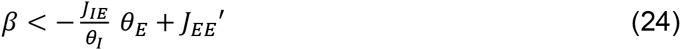

In the intersection of these two semi-planes both D and U are stable (bistable region, Fig. 5A red). In contrast, in the complementary region to the two semi-planes, neither U nor D are stable (Fig. 5A white region). There, a rhythmic concatenation of relatively long U and D periods is observed where the network stays transiently in each state until adaptation triggers a transition (see e.g. Fig. 4E). Because of the separation of time-scales, we refer to this stability to the rate dynamics but not to the adaptation dynamics as *quasi-stable* states.

The addition of noise makes that some of the stable solutions now become meta-stable, meaning that the network can switch to a different state by the action of the noise (i.e. the external fluctuations in our model). This is the case of the bistable region (Fig. 5A red) where fluctuations generate stochastic transitions between the two metastable U and D states (Fig. 4D). In the region of D stability *θ*_*E*_ > 0, we find a new subregion with noise-driven transitions from a metastable D state to a *quasi-stable U state*, and back to D by the action of adaptation (Fig. 5A light violet). This subregion is delimited by the condition that U is not stable (i.e. Eq. 24 does not hold) but *just* because of the existence of adaptation. This can be written by saying that Eq. 24 holds if β=0:

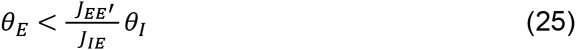

Equivalently, within the region of U stability, noise creates a new subregion with noise-driven transitions from a metastable U state to a *quasi-stable D state*, and back to U by the recovery from adaptation (Fig. 5A light orange). This subregion is given by the condition that there is a negative effective threshold *θ*_*E*_ < 0 (i.e. caused by a supra-threshold mean external drive) but the adaptation *a*^*U*^ recruited in the U state is sufficient to counterbalance it: *a*^*U*^ + *θ*_*E*_ > 0. This makes the D transiently stable until adaptation decays back to zero. Substituting 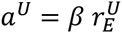 (Eq. 14) and 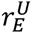 by the equilibrium rate at the U state given by Eq. 18, the limit of this subregion can be expressed as (Fig. 5A, light orange region):

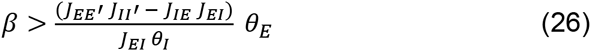

## Acknowledgments

We thank Zhe Chen for sharing the HMM based UP-DOWN detection related scripts, Narani van Laarhoven for sharing codes, Kenneth Harris and the members of the Compte and de la Rocha labs for helpful discussions. This work was supported by the AGAUR of the Generalitat de Catalunya (Ref: SGR14-1265), the Spanish Ministry of Economy and Competitiveness together with the European Regional Development Fund (grants BFU2009-09537, BFU2012-34838 to A.C., RYC-2011-08755 to A.R., SAF2010-15730, SAF2013-46717-R and RYC-2009-04829 to J.R.), the EU (Marie Curie grants PIRG07-GA-2010-268382 to J.R. and BIOTRACK contract PCOFUND-GA-2008-229673 to A.R.). Part of the work was carried out at the Esther Koplowitz Centre, Barcelona.

